# First complete genome sequences of *Streptococcus pyogenes* NCTC 8198^T^ and CCUG 4207^T^, the type strain of the type species of the genus *Streptococcus*: 100% match in length and sequence identity between PacBio solo and Illumina plus Oxford Nanopore hybrid assemblies

**DOI:** 10.1101/2020.03.10.985267

**Authors:** Francisco Salvà-Serra, Daniel Jaén-Luchoro, Hedvig E. Jakobsson, Lucia Gonzales-Siles, Roger Karlsson, Antonio Busquets, Margarita Gomila, Antonio Bennasar-Figueras, Julie E. Russell, Mohammed Abbas Fazal, Sarah Alexander, Edward R. B. Moore

## Abstract

We present the first complete, closed genome sequences of *Streptococcus pyogenes* strains NCTC 8198^T^ and CCUG 4207^T^, the type strain of the type species of the genus *Streptococcus* and an important human pathogen that causes a wide range of infectious diseases. *S. pyogenes* NCTC 8198^T^ and CCUG 4207^T^ are derived from deposit of the same strain at two different culture collections. NCTC 8198^T^ was sequenced, using a PacBio platform; the genome sequence was assembled *de novo*, using HGAP. CCUG 4207^T^ was sequenced and a *de novo* hybrid assembly was generated, using SPAdes, combining Illumina and Oxford Nanopore sequence reads. Both strategies, yielded closed genome sequences of 1,914,862 bp, identical in length and sequence identity. Combining short-read Illumina and long-read Oxford Nanopore sequence data circumvented the expected error rate of the nanopore sequencing technology, producing a genome sequence indistinguishable to the one determined with PacBio. Sequence analyses revealed five prophage regions, a CRISPR-Cas system, numerous virulence factors and no relevant antibiotic resistance genes.

These two complete genome sequences of the type strain of *S. pyogenes* will effectively serve as valuable taxonomic and genomic references for infectious disease diagnostics, as well as references for future studies and applications within the genus *Streptococcus*.

## Introduction

*Streptococcus pyogenes*, within the β-hemolytic, Lancefield group A *Streptococcus* (GAS) (Lancefield, 1933), is an important clinically-relevant and strictly-human pathogen causing a wide range of diseases, including local and invasive infections (e.g., throat, skin infections, meningitis), severe toxin-mediated diseases (e.g., necrotizing fasciitis, scarlet fever, streptococcal toxic shock syndrome) and immune-mediated diseases (e.g., rheumatic fever, rheumatic heart disease, post-streptococcal glomerulonephritis) (Ralph and Carapetis, 2013).

In 2005, it was estimated that more than 500,000 people were dying every year from severe diseases caused by GAS, as well as an estimated 600 million new cases of pharyngitis and 100 million new cases of pyoderma (Carapetis et al., 2005). Thus, *S. pyogenes* is among the top-10 infectious causes of mortality in humans (Barnett et al., 2019). Moreover, *S. pyogenes* is the type species of the genus *Streptococcus*, the type genus of the family *Streptococcaceae*, and as a clinically-relevant bacterium, *S. pyogenes* has been continuously studied since it was first described (Rosenbach, 1884).

In recent decades, several next-generation and third-generation (i.e., long-read) sequencing technologies have emerged and are now widely used in many settings (Loman and Pallen, 2015). For instance, Illumina has led the field in high-throughput DNA sequencing, by providing highly accurate and relatively inexpensive sequence reads. However, their short lengths (few hundred base-pairs) have restricted efficacy to resolve problematic genomic regions (e.g., repeats, ribosomal operons, long sequence motifs), sometimes yielding fragmented and incomplete assemblies (Goodwin et al., 2016). Meanwhile, PacBio provides long reads (several kilobase-pairs) with high consensus accuracy; generally yielding complete bacterial genome sequences. However, high capital costs of PacBio platforms have constrained accessibility to users, who normally access them via commercial/institutional sequencing services. Additionally, requirements of large quantities of high-quality DNA make PacBio sequencing relatively laborious, time-consuming and impractical for some applications. More recently, Oxford Nanopore Technologies launched the MinION portable sequencer, which provides ultra-long reads of as many as two million base-pairs (Payne et al., 2018), requiring simple, rapid and cost-effective DNA library preparation protocols. Nanopore-sequencing has been demonstrated to resolve very-long repetitive regions that not even PacBio-sequencing could resolve (Schmid et al., 2018). However, inaugural high error rates (>30%; currently ~7%) (Greninger et al., 2015, Laver et al., 2015, Goodwin et al., 2015) caused some degree of doubt within the scientific community, although more recent developments and studies have allayed much of the initial skepticism.

Resulting from these technological developments, in 2019-06-29, 1,883 genome sequences of *S. pyogenes* were publicly available in GenBank, of which 195 were complete. However, of those 195, only the complete genome sequences presented in this study represented the type and most important reference strain of the species.

Here, we present the first complete genome sequences of the type strain of *S. pyogenes* (NCTC 8198^T^ = CCUG 4207^T^), determined by two different approaches: *S. pyogenes* NCTC 8198^T^ completed using only PacBio reads; and *S. pyogenes* CCUG 4207^T^ completed by combining Illumina and Oxford Nanopore reads. Both assemblies were identical in length and sequence identity, demonstrating the possibility of surpassing the inherent error rate of the nanopore sequencing technology, by combining Illumina reads and, thus, obtaining an assembly as accurate as the one obtained with the PacBio approach.

## Materials and methods

### Cultivation conditions and DNA extractions

*S. pyogenes* NCTC 8198^T^ was cultivated at the laboratories of the National Collection of Type Cultures (NCTC, London, UK) on Columbia blood agar (Columbia Agar Base plus 5% horse blood, Thermo Fisher Scientific, Waltham, MA, USA), at 37°C. High-molecular weight genomic DNA was isolated, using the MasterPure^™^ Gram Positive DNA Purification Kit (Epicentre, Madison, WI, USA), for PacBio sequencing. *S. pyogenes* CCUG 4207^T^ was cultivated at the laboratories of the Culture Collection University of Gothenburg (CCUG, Gothenburg, Sweden) on Chocolate agar medium (Brain Heart Infusion Agar with 10% heat-lysed defibrinated horse-blood, 15% horse-serum and fresh yeast extract, prepared by the Substrate Unit, Department of Clinical Microbiology, Sahlgrenska University Hospital), with 5% CO2, at 37°C. Genomic DNA was extracted from fresh pure biomass, using a Wizard® Genomic DNA Purification Kit (Promega, Madison, WI, USA), for Illumina sequencing, and a modified version (Salvà-Serra et al., 2018) of a previously described protocol (Marmur, 1961), for Oxford Nanopore sequencing (Figure 1).

**Figure 1.**
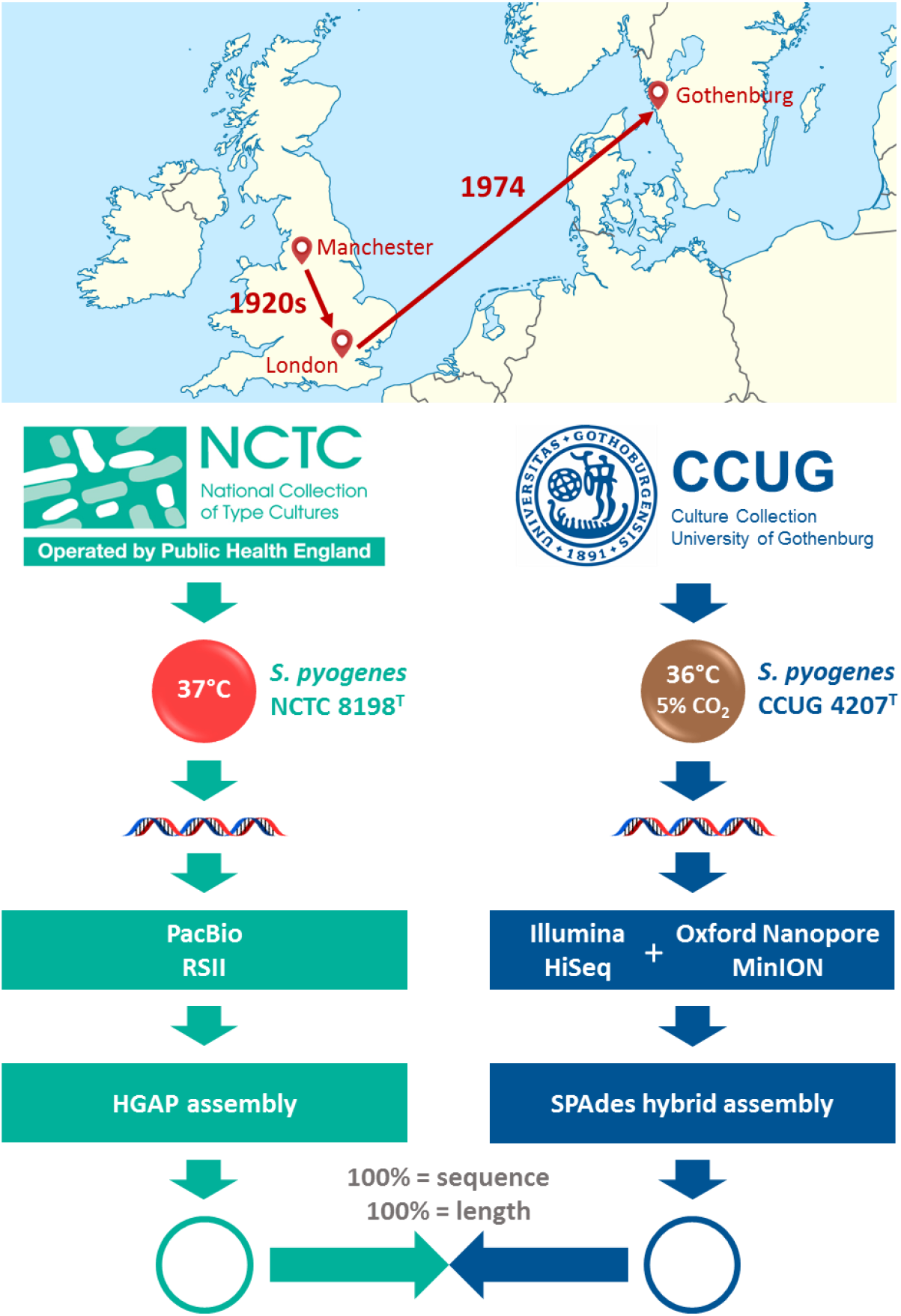
Illustration showing the origin, the strain passage and indicating the whole-genome sequencing workflows performed, in parallel, by the NCTC and the CCUG to determine the sequence of the type strain of *S. pyogenes*.

### PacBio sequencing

High-molecular weight DNA of *S. pyogenes* NCTC 8198^T^ was sheared with a 26 G blunt Luer-Lok^TM^ needle and used to prepare two 10 to 20-kb PacBio SMRT libraries, following the manufacturer’s recommendations. The libraries were sequenced using the P6-C4 chemistry on a Single Molecule, Real-Time (SMRT) cell, using a PacBio RSII platform (Pacific Biosciences of California, Inc., Menlo Park, CA, USA) (www.pacb.com), at the Wellcome Trust Sanger Institute (Hinxton, UK).

### Illumina sequencing

Genomic DNA of *S. pyogenes* CCUG 4207^T^ was used to prepare a standard Illumina library, with an insert size ranging from 130 to 680 bp, following an optimized protocol (GATC Biotech, Konstanz, Germany) and using standard Illumina adapter sequences. The library was sequenced at GATC Biotech (Konstanz, Germany), using an Illumina HiSeq 2500 instrument (Illumina, Inc., San Diego, CA, USA) (www.illumina.com) to generate paired-end reads of 126 bp.

### Oxford Nanopore sequencing

High-molecular weight DNA of *S. pyogenes* CCUG 4207^T^ was used to prepare a sequencing library, using a Rapid Sequencing Kit (SQK-RAD003) (Oxford Nanopore Technologies, Ltd., Oxford, UK), according to manufacturer’s instructions. The library was sequenced at the CCUG laboratories, on a MinION sequencer (Oxford Nanopore Technologies, Ltd., Oxford, UK) (www.nanoporetech.com), using a Flow Cell model FLO-MIN 106 version R9.4. The sequencing was performed with the software MinKNOWN, version 1.10.23, selecting the 48-h sequencing script (*NC_48Hr_sequencing_FLO-MIN106_SQK-RAD003*), with default parameters.

### PacBio *de novo* assembly

PacBio sequence reads from both SMRT sequencing runs were used in the assembly. Read quality was assessed, using NanoPlot version 1.13.0 (De Coster et al., 2018). Sequence reads were auto-error-corrected and assembled *de novo*, using the Hierarchical Genome Assembly Process (HGAP) version 3 (Chin et al., 2013). The assembled sequence was polished with the consensus calling algorithm, Quiver, version 1. The ends of the final assembly were trimmed (i.e., eliminating sequence redundancy) manually, to circularize the genome, and the genome sequence reorganized to start with the *dnaA* gene, which encodes the chromosomal replication initiator protein, DnaA. Assembly statistics were obtained, using QUAST, version 4.5 (Gurevich et al., 2013).

### Illumina-Nanopore hybrid *de novo* assembly

The quality of the Illumina paired-end reads was analyzed with FastQC, version 0.11.3 (https://www.bioinformatics.babraham.ac.uk/projects/fastqc/). Subsequently, the reads were sub-sampled and trimmed, using Sickle, version 1.33 (Joshi and Fass, 2011), with a Phred quality score threshold of Q30. Meanwhile, the FAST5 files containing the raw data generated by the Oxford Nanopore sequencing run were processed with the Oxford Nanopore basecalling pipeline, Albacore, version 2.0.2., and the quality was analyzed with NanoPlot version 1.13.0 (De Coster et al., 2018). Only reads with a quality score great than Q7 were used for the assembly (i.e., classified as *Pass* by Albacore). Afterwards, a hybrid *de novo* assembly, using both Illumina and Oxford Nanopore reads was performed with SPAdes, version 3.11.0 (Bankevich et al., 2012, Antipov et al., 2016). The assembly was performed with the flag --*careful* enabled, to map the Illumina reads back to the assembly with BWA, version 0.7.12-r1039 (Li and Durbin, 2009) and to reduce the number of mismatches and short indels. The ends of the assembly were trimmed manually, and the sequence reorganized to start with *dnaA*, as was done for the PacBio assembly. Assembly statistics were obtained, using QUAST, version 4.5 (Gurevich et al., 2013).

### Genome sequence comparisons

Once assembled, closed and completed, the genome sequences of *S. pyogenes* NCTC 8198^T^ and *S. pyogenes* CCUG 4207^T^ were compared (Figure 1). Firstly, both genome assemblies were aligned, using BLASTN, version 2.2.10 (Altschul et al., 1990). Secondly, all the raw Illumina paired-end reads were mapped against complete and closed genome sequences, using CLC Genomics Workbench, version 10.0 (Qiagen Aarhus A/S, Aarhus, Denmark), and a Basic Variant Detection 2.0 analysis was performed, using the same software, using a minimum frequency of 35% (default).

### Genome annotations and characterization

The genome sequence of *S. pyogenes* NCTC 8198^T^ was initially annotated with Prokka (Seemann, 2014), and submitted to the European Nucleotide Archive (Harrison et al., 2019). Afterwards, the genome sequence was re-annotated, with the NCBI Prokaryotic Genome Annotation Pipeline (PGAP), version 4.1 (Tatusova et al., 2016), and deposited in the NCBI Reference Sequence (RefSeq) database (O’Leary et al., 2016). The genome sequence of *S. pyogenes* CCUG 4207^T^ was submitted to GenBank (Sayers et al., 2019). Subsequently, the sequence was annotated with PGAP, version 4.7, and deposited in RefSeq. The latest of these annotations (i.e., PGAP version 4.7) was used for further analyses and to construct a genome atlas with the on-line server Gview (Petkau et al., 2010).

The on-line tool, PHASTER (Arndt et al., 2016), was used to search for prophages inserted in the chromosome, while the tool CRISPRFinder (Grissa et al., 2007b) was used to search clustered, regularly interspaced short palindromic repeat (CRISPR) arrays. The consensus sequence of the direct repeats were classified, using CRISPRmap v2.1.3-2014 (Lange et al., 2013, Alkhnbashi et al., 2014), and the crRNA-encoding strand determined, using CRISPRstrand (Alkhnbashi et al., 2014), implemented in CRISPRmap. Additionally, the tool CRISPRone (Zhang and Ye, 2017) was used to confirm the detected CRISPR arrays and to identify possible CRISPR-associated genes (*cas*). Spacer sequences of CRISPR arrays were analyzed with BLASTN, version 2.2.10 (Altschul et al., 1990), against the complete genome sequence of *S. pyogenes* NCTC 8198^T^ (= CCUG 4207^T^). Searches for virulence factors across the genome was done against the protein sequences of the curated core dataset (3,200 protein sequences) of the Virulence Factors Database (VFDB; http://www.mgc.ac.cn/VFs/) (Liu et al., 2019), using BLASTP, version 2.2.10 (Altschul et al., 1990). Only hits with ≥50% of identity over ≥50% of the DNA sequence length were considered. For subtyping, the *emm* gene sequence was analyzed, with BLASTN 2.2.27+, against the *emm* gene database of the *Streptococcus* Laboratory (Centers for Disease Control and Prevention, CDC, USA; https://www2a.cdc.gov/ncidod/biotech/strepblast.asp). Potential antibiotic resistance genes were searched with the tool, Resistance Gene Identifier (RGI), of the Comprehensive Antibiotic Resistance Database (CARD; https://card.mcmaster.ca/) (Jia et al., 2017). Only results classified as “perfect” or “strict” were considered.

### Data availability

The complete genome sequence of *S. pyogenes* NCTC 8198^T^ is publicly available in DDBJ/ENA/GenBank under the accession number LN831034. The version described in this article is LN831034.1. The PacBio sequence reads of *S. pyogenes* NCTC 8198^T^ are available in the Sequence Read Archive (SRA) (Leinonen et al., 2011) under the accession numbers ERR550482 and ERR550487. The complete genome sequence of *S. pyogenes* CCUG 4207^T^ is publicly available in DDBJ/ENA/GenBank under the accession number CP028841. The version described in this article is CP028841.1. The Illumina and Oxford Nanopore raw sequence reads of *S. pyogenes* CCUG 4207^T^ are available in the SRA under the accession numbers SRR8631872 and SRR10092043, respectively. The Illumina and Oxford Nanopore sequence reads sets used in the assembly are available in the SRA under the accession numbers SRR10092042 and SRR8608127.

## Results

### Whole-genome sequencing

The complete genome sequence of the type strain of *S. pyogenes* (NCTC 8198^T^ = CCUG 4207^T^) was determined, using three different sequencing technologies. The yield of each sequencing run is summarized in Table 1.

**Table 1.** Results of whole-genome sequencing of *S. pyogenes* NCTC 8198^T^ and CCUG 4207^T^, from the four sequencing runs done with PacBio RSII, Illumina HiSeq 2500 and Oxford Nanopore MinION platforms. The total number of reads, total yield (Mb), sequencing depth, average read length (bp), longest read (bp) and average Phred quality score are shown for each sequencing run. The SRA accession number of each run is indicated.

| Platform | PacBio RSII |  | Illumina HiSeq<br>2500 | Oxford Nanopore<br>MinION |
| --- | --- | --- | --- | --- |
| Run SRA accession<br>number | ERR550482 | ERR550487 | SRR8631872 | SRR10092043 |
| Total number of reads | 163,469 | 163,274 | 10,260,942 | 135,020 |
| Total yield (Mb) | 420 | 694 | 1,293 | 1,122 |
| Sequencing depth | 219 X | 362 X | 675 X | 586 X |
| Average read length (bp) | 2,570 | 4,250 | 126 | 8,307 |
| Read length N50 (bp) | 4,555 | 9,809 | 126 | 14,598 |
| Longest read (bp) | 35,267 | 41,160 | 126 | 97,527 |
| Average Phred quality score | 2.7 | 3.8 | 35 | 11.5 |

#### PacBio

The two PacBio sequencing runs yielded a total of 420 and 694 Mb of raw data, distributed in 163,469 and 163,274 sequence reads, with average lengths of 2,570 and 4,250 bp, respectively. The sequence read length N50s were 4,555 and 9,809 bp, with longest reads of 35,267 and 41,160 bp and mean read Phred quality scores of 2.7 and 3.8, respectively.

#### Illumina

The Illumina genome sequencing of *S. pyogenes* CCUG 4207^T^ yielded 1,292.9 Mb distributed in 10,260,942 paired-end reads of 126 bp with 94.1% of the reads exhibiting an average Phred quality score (i.e., for each sequence read, the arithmetic mean of every base quality) ≥ Q30. The total average Phred quality score was 35. Down-sampling and trimming with Sickle left 355.4 Mb distributed in 3,083,010 paired-end reads of an average length of 115 bp.

#### Oxford Nanopore

The Oxford Nanopore whole-genome sequencing of *S. pyogenes* CCUG 4207^T^ was done, using the Rapid Sequencing Kit SQK-RAD003 with a MinION sequencer, on a Flow Cell (FLO-MIN 106, R9.4) with 465 active channels. The run yielded 1,121.6 Mb distributed in 135,020 reads with a mean length of 8,307 bp; the read length N50 was 14,598 bp and the longest read was 97,527 bp with a mean read Phred quality score of 11.5. In total, 130,764 reads (96.85%) were classified as *Pass* by Albacore and used for the hybrid assembly.

### Genome sequence assemblies

The assembly of the PacBio reads with HGAP, followed by a polishing step performed with Quiver, yielded a complete and closed sequence. Trimming of the ends (i.e., overlapping redundant sequences) resulted in a final sequence of 1,914,862 bp, representing the genome of *S. pyogenes* NCTC 8198^T^. In parallel, the *de novo* hybrid assembly of the trimmed Illumina sequence reads plus the basecalled Oxford Nanopore reads also resulted in a complete and closed sequence. The trimming of the ends yielded a final sequence of 1,914,862 bp. Analysis performed with QUAST confirmed that both assemblies did not contain any gaps (i.e., no N’s).

The final complete and closed genome sequence of *S. pyogenes* CCUG 4207^T^ was analyzed, using BLASTN, against the complete genome sequence of *S. pyogenes* NCTC 8198^T^. The analysis yielded a match of 1,914,862 bp with 100% of identity. Afterwards, for further quality control, the entire set of raw Illumina paired-end reads (i.e., 1,292.9 Mb, coverage: 675 X) was mapped against the two complete genome sequences. A variant calling analysis was performed, and no variants were found in any of the cases. Thus, the two independent and parallel strategies of sequencing and assembly resulted in a genome sequence of identical length and identity of 1,914,862 bp and a GC content of 38.5% (Table 2).

**Table 2.** General information and genomic features of *S. pyogenes* NCTC 8198^T^ = CCUG 4207^T^.

| Section | Features |  |  |
| --- | --- | --- | --- |
| General information | Organism | Streptococcus pyogenes |  |
|  | Taxonomy ID | 1314 |  |
|  | Strain | S.F. 130<sup>T</sup> = NCTC 8198<sup>T</sup> = CCUG 4207<sup>T</sup> |  |
|  | Collection date | ca., 1926 |  |
|  | Host | Homo sapiens |  |
|  | Host disease | Scarlett fever |  |
|  | Geographic location | United Kingdom, Manchester |  |
| Assemblies | Strain | S. pyogenes NCTC 8198<sup>T</sup> | S. pyogenes CCUG 4207<sup>T</sup> |
|  | Sequencing platforms | PacBio RSII | Illumina HiSeq 2500 + Oxford Nanopore MinION |
|  | Assembly method | HGAP version 3 | SPAdes version 3.11.0 |
|  | Assembly coverage | 581 X | 186 X (Illumina) + 576 X (Oxford Nanopore) |
|  | GenBank accession number | LN831034 | CP028841 |
|  | Assembly accession number | GCA_002055535.1 | GCA_004028355.1 |
|  | BioProject ID | NCTC_3000 (PRJEB6403) | TAILORED-Treatment (PRJNA302716) |
|  | Finishing quality | Closed complete genome | Closed complete genome |
|  | Number of contigs | 1 | 1 |
|  | Number of N's | 0 | 0 |
|  | Total length (bp) | 1,914,862 | 1,914,862 |
|  | GC content (%) | 38.5 | 38.5 |
| RefSeq annotation | Annotation method | PGAP version 4.1 | PGAP version 4.7 |
|  | Number of genes (total) | 2,007 | 2,009 |
|  | Total coding sequences (CDSs) | 1,918 | 1,920 |
|  | Protein coding sequences | 1,866 | 1,860 |
|  | Pseudogenes | 52 | 60 |
|  | Number of RNA genes | 89 | 89 |
|  | tRNA | 67 | 67 |
|  | Non-coding RNA | 4 | 4 |
|  | Ribosomal RNA | 18 (6 operons) | 18 (6 operons) |
|  | Hypothetical proteins | 334 | 306 |

### Characterization of the complete genome sequence

#### Annotation

The latest annotation (i.e., *S. pyogenes* CCUG 4207^T^ annotated with PGAP version 4.7 and available in RefSeq) revealed a total of 2,009 genes, of which 1,920 were CDSs. The annotation detected 89 RNA genes, which included 67 tRNA genes, four non-coding RNA genes and 18 ribosomal genes distributed in six complete ribosomal operons. Additionally, 60 pseudogenes were annotated. A total of 306 genes (14.8%) were annotated as “hypothetical proteins”. A genome atlas of this annotation version is depicted in Figure 2.

**Figure 2.**
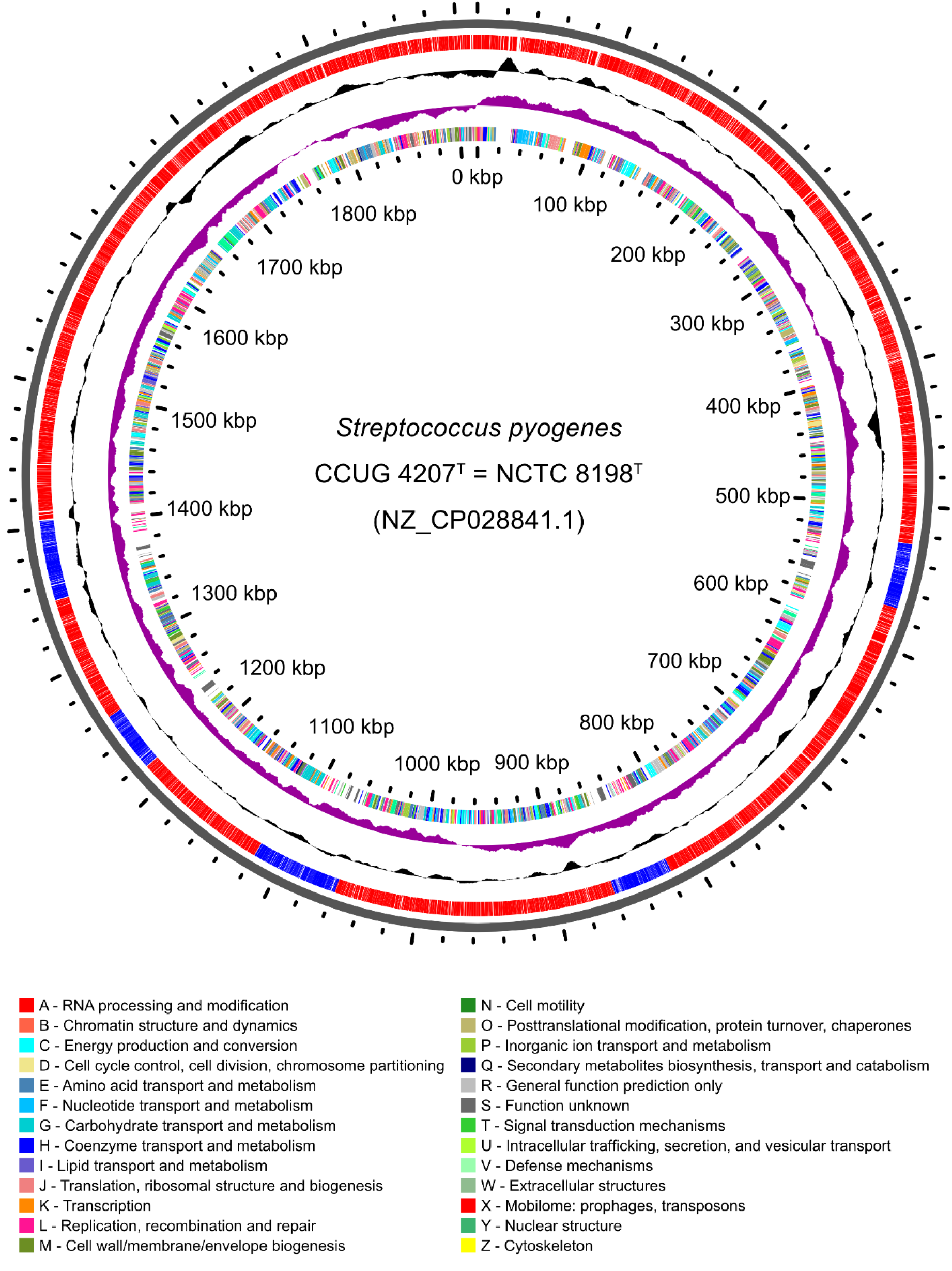
Genome atlas of the type strain of *Streptococcus pyogenes*. The atlas was built with the genome sequence annotated with PGAP 4.7 and available in RefSeq (NZ_CP028841.1). Labeling, from outside to inside: backbone; CDSs and prophage regions (colored in blue); GC content; GC skew and CDSs colored by COG categories (if assigned).

#### Prophages

Five putative prophages were detected using the software PHASTER, four marked by the software as ‘intact’ (completeness score > 90) and one as ‘questionable’ (completeness score = 70-90), with a GC content ranging from 37.4 to 39.1% (Table 3). The largest prophage region was 56,926 bp and the shortest 41,886 bp. Overall, the five regions add up to 234,671 bp, which represents 12.3% of the genome size (Figure 2). In total, according to the PGAP annotation, the five prophage regions encompass 321 CDSs, which represents a 16.72% of the total 1,920 CDSs annotated by PGAP in the genome sequence.

**Table 3.** Prophage regions identified by the software PHASTER. For each region, the estimated completeness, the positions in the genome, length, number of CDSs and percentage of GC are indicated.

| Region | Completeness | Start | End | Length (bp) | No. CDSs | GC content (%) |
| --- | --- | --- | --- | --- | --- | --- |
| **SF130.1** | Questionable | 526,644 | 570,537 | 43,894 | 65 | 38.6 |
| **SF130.2** | Intact | 818,527 | 860,412 | 41,886 | 59 | 38.3 |
| **SF130.3** | Intact | 1,058,125 | 1,107,536 | 49,412 | 67 | 37.4 |
| **SF130.4** | Intact | 1,216,850 | 1,259,402 | 42,553 | 60 | 37.7 |
| **SF130.5** | Intact | 1,348,482 | 1,405,407 | 56,926 | 70 | 39.1 |

#### CRISPR-Cas systems

The analysis of the genome with CRISPRFinder revealed the presence of a CRISPR array (positions: 1,317,644 – 1,317,938 bp), composed of five direct repeats of 32 bp and four spacers of sizes ranging from 33 to 35 bp. The consensus sequence of the direct repeats was classified, with CRISPRmap, into the family 5 and structure motif 3 (Figure 3). The analysis with CRISPRone revealed seven CRISPR-associated (*cas*) genes, located adjacent to the CRISPR array (*cas3*, *cas5*, *cas8c*, *cas7*, *cas4*, *cas1* and *cas2*; locus tags: DB248_RS07080 - DB248_RS07050). This architecture corresponds to the Class 1, subtype I-C of the updated evolutionary classification of CRISPR-Cas systems (Makarova et al., 2015). However, *cas3* was frame-shifted due to a single-nucleotide deletion. The frameshift was confirmed by manually inspecting the mapped Illumina reads.

**Figure 3.**
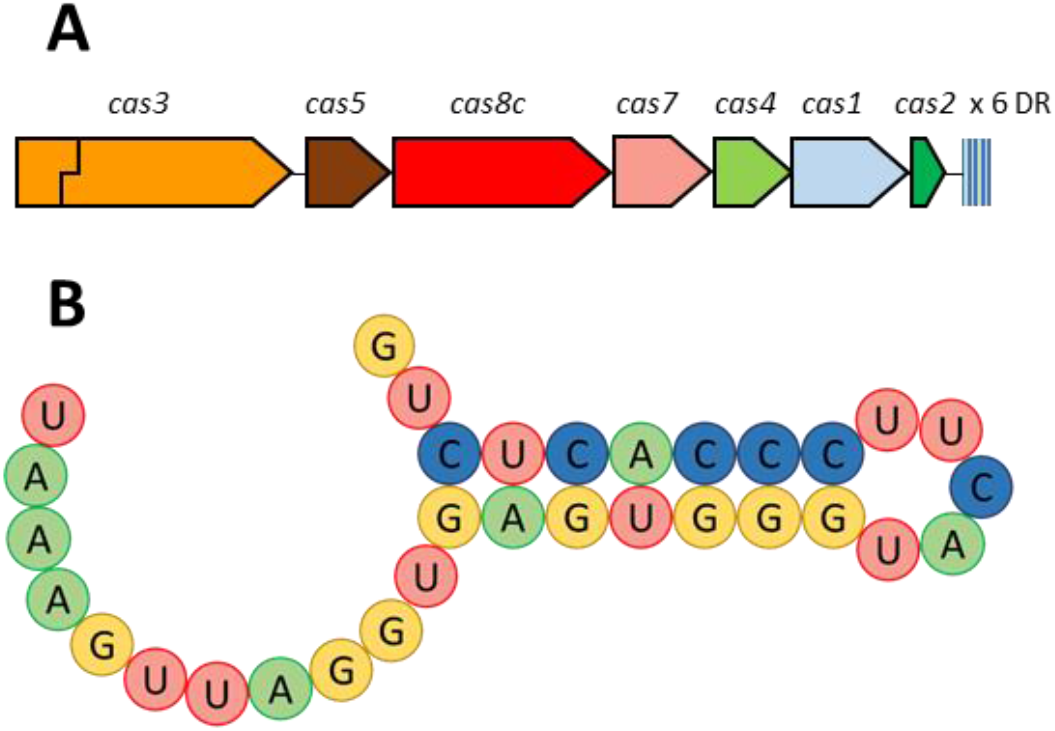
(A) The genomic architecture of the *S. pyogenes* CCUG 4207^T^ (= NCTC 8198^T^) CRISPR-Cas subtype I-C system, formed by seven *cas* genes followed by six direct repeats (DR) and five spacers. The stepped line crossing the *cas3* gene indicates frameshift. (B) The hairpin structure motif of the consensus sequence of the direct repeats of the CRISPR array of *S. pyogenes* CCUG 4207^T^ (= NCTC 8198^T^).

The BLASTN analyses of the spacer sequences against the whole genome sequence revealed that the sequence of the second spacer of the CRISPR array (positions 1,317,807 – 1,317,839 bp) shows 100% identity against a sequence of the prophage region SF130.1 (positions: 551,171 – 551,203; identities = 33/33), located in a gene encoding a phage predominant capsid protein (DB248_RS03050). Additionally, the sequence of the third spacer shows high similarity to a sequence of the prophage region SF130.5 (positions: 1,388,239 – 1,388,206 bp; identities: 32/34), which is part of a gene encoding a hypothetical protein (DB248_RS07380). Finally, the sequence of the fourth spacer presents a high degree of homology to a sequence of the prophage region SF130.2 (positions: 850,316 – 850,348; identities = 32/33), which is also part of a gene encoding a hypothetical protein (DB248_RS04545).

Additionally, four putative *cas* genes (*cas9*, *cas1*, *cas2*, *csn2*; locus tags: DB248_RS04305 - DB248_RS04320) were located between positions 812,387 and 818,352, adjacent to a gene encoding a phage integrase of the prophage region 2 (DB248_RS04325). The architecture of this locus corresponds to the Class 2, subtype II-A of the CRISPR-Cas systems classification, although no CRISPR arrays were found in the surrounding region.

#### Virulence factors

The protein sequences of *S. pyogenes* CCUG 4207^T^ (= NCTC 8198^T^) were analyzed with BLASTP against the core dataset of the Virulence Factors Database VFDB. Numerous genes encoding putative virulence factors were detected (Table S1), several of them within the prophage regions. One of the genes was *emm*, encoding the surface protein M (DB248_RS09295), one of the major virulence factors of *S. pyogenes*, which provides protection against the immune system and has been used for strain serological typing (Metzgar and Zampolli, 2011). The BLATN analysis of the sequence of the *emm* gene against the *emm* database of the *Streptococcus* Laboratory (CDC, USA) confirmed that it is type 1.0. Genes related with the synthesis of the hyaluronic acid capsule were also found (*hasA*, DB248_RS09950; *hasB*, DB248_RS09955; *hasC*, DB248_RS09965). This polysaccharide capsule is a key virulence factor involved in adhesion, tissue invasion (Cywes and Wessels, 2001) as well as in molecular mimicry for immune evasion (Wessels et al., 1991). The *sic* gene encoding the hypervariable streptococcal inhibitor of complement-mediated lysis (SIC) was also found (DB248_RS09285) (Fernie-King et al., 2001). Two genes encoding DNAses were also found: *mf/spd* (DB248_RS09395), encoding the DNaseB (Sriskandan et al., 2000, Iwasaki et al., 1997) and *mf3/spd3* (DB248_RS06505), within the prophage region SF130.4, encoding the DNaseC (Sumby et al., 2005). A gene encoding the hyaluronate lyase HylA was detected (DB248_RS04240). HylA has been suggested to facilitate spread of large molecules and to play a nutritional role for *S. pyogenes*, by disrupting host-tissue as well as its own capsule, allowing growth on hyaluronic acid as carbon source (Starr and Engleberg, 2006). In addition, four prophage-associated hyaluronidase-encoding genes were found (DB248_RS03100, DB248_RS04540, DB248_RS06545, DB248_RS07275), located in the prophage regions SF130.1, SF130.2, SF130.4 and SF130.5, respectively, which may act as additional spreading factors (Hynes, 2004). These enzymes seem to be useful for phages to penetrate the capsule of hyaluronic acid (Smith et al., 2005). Additionally, an *ideS/mac* gene, encoding an immunoglobulin G-degrading enzyme, was found (DB248_RS03795). This exoenzyme shields the cells from being opsonized by IgG antibodies, by cleaving their heavy chain (von Pawel-Rammingen et al., 2002). A gene encoding a C5a peptidase was also found (*scpA*, DB248_RS09275). This peptidase degrades the chemotactic complement factor C5a (Cleary et al., 1992), thus preventing the C5a-based recruitment of neutrophils and other inflammatory cells to the site of infection (Guo and Ward, 2005). Moreover, a gene encoding the streptokinase A (*ska*, DB248_RS09135) was found, which catalyzes the conversion of plasminogen to plasmin, a serine protease that facilitates tissue invasion by degrading proteins of the extracellular matrix (Ringdahl et al., 1998). Additionally, several genes encoding putative streptococcal exotoxins were detected: *speA* (DB248_RS05660), encoding a pyrogenic exotoxin type A and located in the prophage region SF130.3; *speB* (DB248_RS09375), encoding a pyrogenic exotoxin type B; *speG* (DB248_RS01135), encoding a pyrogenic exotoxin type G; *speJ* (DB248_RS01990) encoding a pyrogenic exotoxin type J precursor; and *smeZ* (DB248_RS09220), encoding the streptococcal mitogenic exotoxin Z. Genes for pili biosynthesis were also detected (*cpa*, DB248_RS00770; *lepA*, DB248_RS00775; *fctA*, DB248_RS00780; *srtC1*, DB248_RS00785; *fctB*, DB248_RS00790), which have been shown to play roles in biofilm formation and attachment to pharyngeal cells (Manetti et al., 2007). Moreover, several genes encoding putative adhesion-related proteins were found (e.g., fibronectin binding proteins), such as *fbaA* (DB248_RS09270), encoding an F2-like fibronectin-binding protein, or *fbp54* (DB248_RS04160). FBP54 has been shown to play a role in the adhesion of GAS to host cells (Courtney et al., 1994, Courtney et al., 1996). An *lbp* gene was also detected (DB248_RS09265), codifying the Lbp laminin-binding protein, involved in adhesion to epithelial cells (Terao et al., 2002) and suggested to play a role in zinc homeostasis (Linke et al., 2009). A *grab* gene, encoding a G-related α2-macroglobulin-binding protein (GRAB), has also been detected, (DB248_RS06200). GRAB is a surface protein that inhibits unwanted proteolysis through a high affinity for α2-macroglobulin, a proteinase inhibitor of human plasma (Rasmussen et al., 1999).

#### Antibiotic resistances

The analysis of the genome sequences with RGI (CARD) revealed one gene related with antibiotic resistance and classified as “strict”. The gene encodes a putative ABC transporter ATP-binding protein (locus tag: DB248_01185), located downstream of a gene encoding another ABC transporter ATP-binding protein (DB248_01180). Gene products show 67 and 66% sequence identity to ABC transporters PatB and PatA of *S. pneumoniae* TIGR4, with overexpression linked to fluoroquinolone resistance (Garvey et al., 2011).

## Discussion

Here we present the first complete genome sequence of *S. pyogenes* NCTC 8198^T^ = CCUG 4207^T^, the type strain of the type species of the genus *Streptococcus*, the type genus of the family *Streptococcaceae*. The sequence has been determined twice, using two fully independent but parallel strategies: PacBio-solo and Illumina plus Oxford Nanopore sequencing. Both strategies have yielded 100% identical complete genome sequences, thus demonstrating that hybrid approaches can completely mitigate the error rate of long read sequences.

The type strain of *S. pyogenes* (NCTC 8198^T^ = CCUG 4207^T^) was isolated as strain S.F. 130 in Manchester, UK, from a throat swab of a scarlet fever case. The strain was provided by William C. C. Topley (University of Manchester) to Frederick Griffith (Pathological Laboratory of the Ministry of Health), who used it for the preparation of Type 1 agglutination sera, in a study of scarlatinal streptococci (Griffith, 1926). In 1950, the strain was deposited at the NCTC by Robert E. O. Williams (Central Public Health Laboratory, Colindale, London, UK) and, in 1974, the NCTC strain was deposited at the CCUG (Figure 1). After decades being available to the scientific community, the strain has served as a taxonomic reference point and has been used in numerous studies. Today, the strain is also preserved and publicly available in other culture collections, e.g., ATCC 12344^T^ (USA) = BCRC 14758^T^ (Taiwan) = CECT 985^T^ (Spain) = CIP 56.41^T^ (France) = DSM 20565^T^ (Germany) = JCM 5674^T^ (Japan) = LMG 14700^T^ (Belgium) = S.F. 130^T^. To date, nearly 2,000 genome sequences of *S. pyogenes* are available in GenBank/ENA/DDBJ, of which nearly 200 are complete genome sequences. In fact, the first complete genome sequence of a strain of *S. pyogenes* was determined almost twenty years ago (Ferretti et al., 2001). However, despite the clinical relevance of the species and the taxonomic importance of the type strain, this is the first complete genome sequence of the type strain of *S. pyogenes* that has been determined.

Following sequencing, two different sequence assembly strategies were used. While *S. pyogenes* NCTC 8198^T^ was assembled *de novo*, using only PacBio reads, *S. pyogenes* CCUG 4207^T^ was assembled *de novo*, using both short Illumina and long Oxford Nanopore reads. Surprisingly, both approaches yielded fully identical complete genome sequences of 1,914,862 bp. Recently, numerous studies have reported high quality genome assemblies, obtained by combining high-quality Illumina reads and long Oxford Nanopore reads (Wick et al., 2017a, De Maio et al., 2019, Goldstein et al., 2019). However, to our knowledge, this is the first study reporting two identical complete genome sequences determined with different methodologies.

The fact that two wholly independent approaches have yielded identical sequences demonstrates the high quality of these genome sequences. On the one hand, the NCTC strain was sequenced, using a PacBio RSII platform, which, as expected, yielded long and relatively inaccurate raw sequence reads (indicated in Table 1, by the low Phred quality scores). However, due to their random distribution, sequencing errors can be corrected during assembly, by high coverage (Koren et al., 2013). On the other hand, the CCUG strain was sequenced with the highly accurate Illumina HiSeq 2500 and the Oxford Nanopore MinION device, which, as expected, yielded long raw sequence reads with low accuracy (indicated in Table 1), Interestingly, Nanopore reads exhibit a higher average Phred quality score than PacBio reads. However, because of the less random distribution of the errors (e.g., misinterpretation of homopolymers) (Lu et al., 2016), the inclusion of the high-quality Illumina reads, during or after the assembly, is crucial to obtain an accurate genome sequence.

These results demonstrate how a hybrid strategy, combining Illumina and Oxford Nanopore sequencing, can provide results as accurate as high coverage PacBio-solo sequencing. This should help to reduce the skepticism generated by the initially high error rate of Oxford Nanopore (Greninger et al., 2015, Laver et al., 2015, Goodwin et al., 2015). In this study, the identical results obtained by both strategies are an indicator of the high quality and accuracy of this genome sequence, which makes it a definitive genomic reference of the species as well as a good model candidate for being used in future evaluations of bacterial genome assemblers.

Furthermore, it is noteworthy that the hybrid assembly was performed with SPAdes (i.e., a relatively user-friendly, well-established and widely-used *de novo* genome assembler), as the assembly itself only required a single command line and did not involve complex and tedious methodologies. Nevertheless, further strategies have been developed in recent years to perform *de novo* hybrid assemblies combining short and long reads (e.g., Unicycler and MaSuRCA), aiming to cover the shortcomings of each technology with the advantages of the other (Wick et al., 2017b, Zimin et al., 2017). Alternative strategies involve only-Nanopore *de novo* assemblies (Koren et al., 2017), which can be afterwards ‘polished’ by other pieces of software (e.g., Pilon and/or Racon) in order to improve their accuracy (Walker et al., 2014, Vaser et al., 2017). In addition, all these strategies and methodologies can be complementary to each other, as one protocol might be more or less useful under particular conditions, while the other one could be the opposite. For instance, SPAdes relies on the short reads to create a first draft assembly and afterwards perform a scaffolding step, while Canu performs an only-long-read *de novo* assembly (Koren et al., 2017) which can be optionally followed by a polishing step with short reads, using software like Pilon (Walker et al., 2014).

In any case, despite the lack of differences between the two determined genome sequences, genomic variations could have been expected, as there have been several passages between strain NCTC 8198^T^ and strain CCUG 4207^T^, and cultivation conditions and DNA preparation methods were different between both culture collections. These circumstances evidently increase the probability of having natural genotypic and phenotypic changes (Somerville et al., 2002, Eberhard et al., 2001, Rezcallah et al., 2004). As a practical example, the ATCC recommends users to do no more than five passages from ATCC^®^ Genuine Cultures. For this reason, i.e., to reduce risk of natural alterations, the same starting material and DNA preparation should had been used. Nonetheless, this genome sequence will be a definitive reference of the type strain of *S. pyogenes*.

The vast amount of genomic data that the current “next-generation” and “third-generation” sequencing platforms generate, can be used for bacterial systematics and taxonomy, which traditionally has been based on observations of phenotypic features, DNA G+C content, DNA-DNA hybridization similarities and sequence determinations and analyses of marker genes, such as 16S rRNA (Konstantinidis and Tiedje, 2005). Recently, numerous studies have shown the effectivity, high resolution and discriminative power of whole-genome sequence-based comparative studies (Gomila et al., 2015, Jensen et al., 2016). In fact, several methods and tools have been developed for analyzing whole-genome sequence similarities, i.e., Average Nucleotide Identity (ANI) (Goris et al., 2007), which can be calculated, using JSpecies (Richter and Rosselló-Móra, 2009, Richter et al., 2016) and *in silico* DNA-DNA hybridization, which can be calculated, using the Genome-to-Genome Distance Calculator (GGDC) (Meier-Kolthoff et al., 2013). Other interesting tools include the Type Strain Genome Server (Meier-Kolthoff and Göker, 2019) and TrueBac^TM^ ID (Ha et al., 2019), both high-throughput on-line servers for genome sequence-based taxonomy, dependent upon curated databases of bacterial species type strain genome sequences. Despite the availability of such tools, public databases contain numerous misclassified genome sequences (Beaz-Hidalgo et al., 2015, Gomila et al., 2015, Jensen et al., 2016), most likely due to disregard for taxonomic controls, but also because genome sequences of the type strains of many species have not yet been determined (Wu et al., 2018) or are not reliable, even because genome sequences may have been erroneously labeled as “type strains” (Salvà-Serra et al., 2019). Global efforts and initiatives are underway to curate public databases (Federhen et al., 2016), as well as to sequence the genomes of type strains (Mukherjee et al., 2017, Wu et al., 2018). Since January 2018, the International Journal of

Systematic and Evolutionary Microbiology (IJSEM), the official publication of the International Committee on Systematics of Prokaryotes (ICSP) and the journal of record for publication of novel microbial taxa, has required authors describing new taxa to provide the genome sequence data; the genome of an organism encodes the basis of its biology and, therefore, is the fundamental basis of information for understanding the organism. Furthermore, genome sequences of the type strains of bacterial and archaeal species are crucial as reference points for identifying and classifying genetic and metagenomic data (Klenk and Göker, 2010, Whitman, 2015).

Five prophage regions have been detected in the genome sequence of *S. pyogenes* CCUG 4207^T^ (= NCTC 8198^T^), encompassing a 17% of the CDSs of the genome. These results agree with initial reports of genome sequences of *S. pyogenes*, which already confirmed a high prevalence of prophages inserted in chromosomes of the species (Ferretti et al., 2001, Beres et al., 2002, Smoot et al., 2002). Overall, numerous studies have shown the crucial role of bacteriophages in the ecology, pathogenicity and the evolution of *S. pyogenes* strains. In fact, prophages have been linked with the recent resurgence of M1-GAS-associated invasive diseases (Nasser et al., 2014).

CRISPR-Cas systems are adaptive immune systems that are widely spread in bacteria (Grissa et al., 2007a, Makarova et al., 2015). These systems have been previously found in several strains of *S. pyogenes*, and an inverse correlation has been observed between them and the number of prophages inserted in the genome (Nozawa et al., 2011). Contrary to this observation, we found a complete CRISPR-Cas system together with five putative prophages in the genome sequence of *S. pyogenes* CCUG 4207^T^ (= NCTC 8198^T^). Additionally, high similarity was found between the sequences of the spacers of the CRISPR arrays and three of the prophage regions, one of them being 100% identical. However, we detected a frameshift, caused by a single-nucleotide deletion in the *cas3* gene, which encodes an endonuclease/helicase that is essential for CRISPR-Cas interference (Brouns et al., 2008). Therefore, the CRISPR-Cas system most likely is non-functional due to this truncation, thus leaving a freeway for infection by bacteriophages, which could explain the presence of five prophage regions inserted within the chromosome. In addition, the short number of spacers also suggests that the CRISPR-Cas system might also not be active in acquisition of new spacers.

As a major human pathogen, *S. pyogenes* has a great repertoire of virulence factors, some of which are intrinsic and shared among almost all strains, while others might be present only in certain strains (Liu et al., 2019) or serotypes (e.g., EndoS2, exclusive of serotype M49) (Sjögren et al., 2013). In this study, the analysis of the protein sequences of *S. pyogenes* CCUG 4207^T^ (= NCTC 8198^T^) against the VFDB has provided insight into the virulence potential of this strain, revealing the presence of numerous prominent virulence factors. In particular, this strain was isolated from a throat swab from a scarlet fever patient (Griffith, 1926). For many years, scarlet fever has been associated with *S. pyogenes* strains producing pyrogenic toxins (Dick and Dick, 1924). In accordance to this, several streptococcal pyrogenic exotoxins have been found, one of them encoded in a prophage region. Additionally, several of the other virulence factors have also been found encoded in prophage regions, highlighting the role of prophages in the pathogenicity of *S. pyogenes* strains. In any case, further studies will be needed to uncover the full pathogenic potential of this strain, with emphasis in revealing the role of the still high number of CDSs annotated as ‘hypothetical proteins’, which represent a 15% of the 2,009 genes annotated.

The only antibiotic resistance-gene detected in the genome sequence was a gene encoding an ABC transporter ATP-binding protein, wherein overexpression has been associated with fluoroquinolone resistance. This lack of significant antibiotic resistance genes was expected, as the type strain of *S. pyogenes* was isolated before the antibiotic era (i.e., before 1928). In addition, the current antibiotic resistance problem is not that great yet among *S. pyogenes* as it is among other bacterial species and taxa, with penicillin remaining the drug of choice, despite numerous decades of use (Spellerberg and Brandt, 2016).

## Conclusions

Here we present the first complete genome sequences of the type strain of *S. pyogenes* (NCTC 8198^T^ = CCUG 4207^T^ = S.F. 130^T^), the type species of the genus *Streptococcus*, the type genus of the family *Streptococcaceae* and a major human pathogen. These genome sequences represent the reference genomic material to be used in taxonomic studies involving this family and its members. Additionally, we have shown how the combination of high-quality, short, Illumina sequence reads with long Oxford Nanopore sequence reads is able to generate a complete genome sequence, identical to the one obtained with only PacBio sequencing.

## Acknowledgements

This work was supported by the European Commission 7^th^ Framework Programme: TAILORED-Treatment (project no. 602860), by the Swedish Västra Götaland regional funding (projects no. ALFGBG-437221 and ALFGBG-720761), the Swedish Västra Götaland FoU grant no. VGFOUREG-665141, Laboratoriemedicin FoU (project N° 51060-6268) and by the Wellcome Trust (project N° 101503/Z/13/Z). DJL, RK and ERBM acknowledge the support of the Joint Programme Initiative - Anti-Microbial Resistance (JPIAMR) (Vetenskapsrådet Project N° 2016-06504). The Culture Collection University of Gothenburg (CCUG) was supported by the Department of Clinical Microbiology, Sahlgrenska University Hospital, Gothenburg, Sweden. FS-S was supported by a stipend for Basic and Advanced Research from the CCUG, through the Institute for Biomedicine, Sahlgrenska Academy, University of Gothenburg. FS-S, DJ-L, HEJ, LG-S, RK, NK and ERBM acknowledge the support from the Centre for Antibiotic Resistance Research (CARe) at the University of Gothenburg. Antonio Busquets was supported by a postdoctoral contract from the University of the Balearic Islands. The authors acknowledge Gemma Langridge, Julian Parkhill and Nick Grayson, at the Wellcome Trust Sanger Institute (Hinxton, United Kingdom), for the PacBio assembly and thank them and Timur Tunovic for valuable comments and suggestions.

## Supplemental material

**Table S1.** Genes encoding putative virulence factors, identified using the Virulence Factors Database.

| VFDB category | Gene | Locus tag | Phage region | Virulence factor |
| --- | --- | --- | --- | --- |
| **Antiphagocytosis** | *emm* | DB248_RS09295 | - | M protein type 1 |
|  | *hasA* | DB248_RS09950 | - | Hyaluronic acid capsule |
|  | *hasB* | DB248_RS09955 | - | Hyaluronic acid capsule |
|  | *hasC* | DB248_RS09965 | - | Hyaluronic acid capsule |
|  | *sic* | DB248_RS09285 | - | SIC (streptococcal inhibitor of complement-mediated lysis) |
| **Exoenzymes** | *mf/spd* | DB248_RS09395 | - | DNaseB |
|  | *mf3/spd3* | DB248_RS06505 | SF130.4 | DNaseC |
|  | *hylA* | DB248_RS04240 | - | Hyaluronidase |
|  | *hylP* | DB248_RS03100 | SF130.1 | Hyaluronidase |
|  | *hylP* | DB248_RS04540 | SF130.2 | Hyaluronidase |
|  | *hylP* | DB248_RS06545 | SF130.4 | Hyaluronidase |
|  | *hylP* | DB248_RS07275 | SF130.5 | Hyaluronidase |
|  | *ideS/mac* | DB248_RS03795 | - | IgG-degrading enzyme |
| **Immune evasion** | *scpA* | DB248_RS09275 | - | C5a peptidase |
| **Plasminogen activator** | *ska* | DB248_RS09135 | - | Streptokinase |
| **Toxin** | *speA* | DB248_RS05660 | SF130.3 | Streptococcal pyrogenic exotoxin A |
|  | *speB* | DB248_RS09375 | - | Streptococcal pyrogenic exotoxin B |
|  | *speG* | DB248_RS01135 | - | Streptococcal pyrogenic exotoxin G |
|  | *speJ* | DB248_RS01990 | - | Streptococcal pyrogenic exotoxin J |
|  | *smeZ* | DB248_RS09220 | - | Streptococcal mitogenic exotoxin Z |
| **Adherence** | *cpa* | DB248_RS00770 | - | Pili |
|  | *lepA* | DB248_RS00775 | - | Pili |
|  | *fctA* | DB248_RS00780 | - | Pili |
|  | *srtC1* | DB248_RS00785 | - | Pili |
|  | *fctB* | DB248_RS00790 | - | Pili |
|  | *fbaA* | DB248_RS09270 | - | FBPs (Fibronectin binding proteins) |
|  | *fbp54* | DB248_RS04160 | - | FBPs (Fibronectin binding proteins) |
|  | *lpb* | DB248_RS09265 | - | Laminin-binding protein |
| **Anti-proteolysis** | *grab* | DB248_RS06200 | - | GRAB (G-related $\alpha_2$-macroglobulin-binding protein) |

